# Estimating the prevalence of missing experiments in a neuroimaging meta-analysis

**DOI:** 10.1101/225425

**Authors:** Pantelis Samartsidis, Silvia Montagna, Angela R. Laird, Peter T. Fox, Timothy D. Johnson, Thomas E. Nichols

## Abstract

Coordinate-based meta-analyses (CBMA) allow researchers to combine the results from multiple fMRI experiments with the goal of obtaining results that are more likely to generalise. However, the interpretation of CBMA findings can be impaired by the file drawer problem, a type of publications bias that refers to experiments that are carried out but are not published. Using foci per contrast count data from the BrainMap database, we propose a zero-truncated modelling approach that allows us to estimate the prevalence of non-significant experiments. We validate our method with simulations and real coordinate data generated from the Human Connectome Project. Application of our method to the data from BrainMap provides evidence for the existence of a file drawer effect, with the rate of missing experiments estimated as at least 6 per 100 reported.

## 1 Introduction

Now over 25 years old, functional magnetic resonance imaging (fMRI) has made significant contributions in improving our understanding of the human brain function. However, the inherent limitations of fMRI experiments have raised concerns regarding the validity and replicability of findings (Farah, 2014). These limitations include poor test-retest reliability (Raemaekers *et al*., 2007), excess of false positive findings (Wager *et al*., 2009) and small sample sizes (Carp, 2012). Meta-analyses play an important role in the field of fMRI as they provide a means to address the aforementioned problems by synthesising the results from multiple experiments and thus draw more reliable conclusions. Since the overwhelming majority of authors rarely share the full data, *coordinate-based meta-analyses* (CBMA), which use the *xyz* coordinates (foci) of peak activations that are typically published, are the main approach for the meta-analysis of fMRI data.

As in any meta-analysis, the first step in a CBMA is a literature search. During this step investigators use databases to retrieve all previous work which is relevant to the question of interest (Normand, 1999). Ideally, this process will yield an exhaustive or at least representative sample of studies on a specific topic. Unfortunately, literature search is subject to the *file drawer* problem (Rosenthal, 1979; Iyengar and Greenhouse, 1988). This problem refers to research studies that are initiated but are not published. When these studies are missing at random (i.e. the reasons that they remain unpublished are independent of their findings), then the pool of studies reduces but the results of a meta-analysis remain reliable. However, if the censoring relates to the findings of a study (e.g. due to rejection by journals reluctant to publish negative results), then meta-analyses may yield biased estimates of the effect of interest (Begg and Berlin, 1988; Sutton *et al*., 2000).

In CBMA, the unit of observation is a *contrast*/*experiment* (these terms are used interchangeably throughout the paper) and not a *study*/*paper* (these terms are also used interchangeably throughout the paper), because the latter may include multiple contrasts that can be used in a single meta-analysis. Hence the file drawer includes contrasts that find no significant activation clusters i.e. ones that report no foci. Such experiments often remain unpublished because when writing a paper, authors focus on the other, significant experiments that they conducted. Moreover, even if mentioned in the final publication, these contrasts are typically not mentioned in the table of foci, and are not registered in the databases which researchers use to retrieve data for their CBMA. The bias introduced by not considering contrasts with no foci depends on how often these occur in practice. For example, if only 1 out of 100 contrasts is null then not considering zero-count contrasts is unlikely to have an impact on the results of a CBMA. However, if this relative proportion is high, then the findings of a CBMA will be misleading in that they will overestimate the effect of interest.

Some authors have attempted to assess the evidence for the existence of publication biases in the field of fMRI. One example is Jennings and Van Horn (2012), who found evidence for publication biases in 74 studies of tasks involving working memory. The authors used the maximum test statistic reported in the frontal lobe as the effect estimate in their statistical tests. Another example is David *et al*. (2013), who studied the relation between sample size and the total number of activations and reached similar conclusions as Jennings and Van Horn (2012). However, to date there has been no work on estimating a fundamental file drawer quantity, that is the prevalence of null experiments.

In this paper, we propose a model for estimating the prevalence of zero-count contrasts in the context of CBMA. Our approach is outlined in Figure 1. Let the *sampling frame* be all *K* neuroimaging experiments of interest that were completed, published or not, where each element of the sampling frame is a statistic map for a contrast. For any contrast in the sampling frame, let *π*(*n*|***θ***) be the probability mass function of the number of foci per contrast, where ***θ*** is a vector of parameters. Hence, null contrasts occur with probability *p*_0_ = *π*(0 | ***θ***) and there are *Kp*_0_ in total. These are unavailable for meta-analysis due to lack of significance. However, the remaining *k* = *K*(1 − *p*_0_) significant experiments are published and are available to draw inference regarding ***θ***. This allows us estimate *p*_0_ and thus the prevalence of zero-count contrasts in the population.

**Figure 1:**
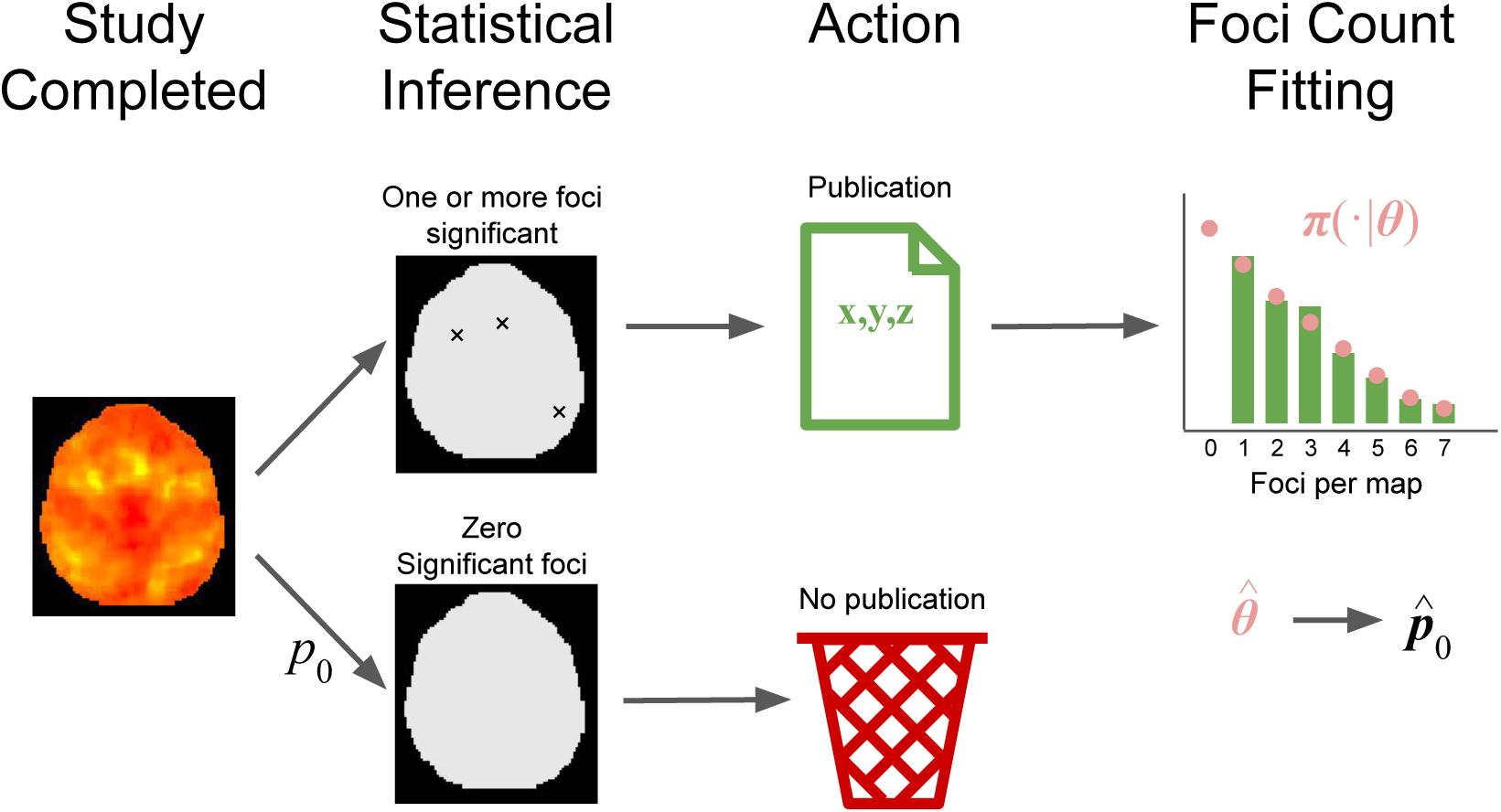
Graphical outline of our approach. We start a population of *K* experiments, where *k* of them are published and the remaining *K* − *k* are not observed due to lack of significance. Assuming *π*(*n*|***θ***) to be the probability mass function of the number of foci per experiment, we use the *k* published experiment to draw inference on ***θ*** and hence estimate *p*_0_ = *π*(0 | ***θ***).

Note that rarely if ever will be able to know the total count of contrasts *k*, no less *K*. Hence our approach can be used to learn the *relative* but not the *absolute* frequency of null contrasts. For the latter, it would be necessary to know *k*, however it is not possible. While our method cannot estimate *p*_0_ in individual meta-analyses, *p*_0_ reflects the sampled population, and thus relates to all studies that make up the population.

Finally, we are careful not to describe our estimators as ‘null study’ prevalence. Rather, we are estimating prevalence of null contrasts. Each study (paper) consists of multiple contrasts, some of which might be null. Therefore, since studies typically involve multiple contrasts, we expect that the prevalence of missing studies is much lower compared to the prevalence of missing experiments.

The remainder of the paper is organised as follows. In Section 2, we describe the CBMA data, both real and simulated, that we used and the statistical model for point data that accounts for missing experiments. In Section 3, we present the results of our simulation studies and real data analyses. Finally, in Section 4 we conclude with a discussion of our main findings and set directions for future research.

## 2 Methods and Materials

### 2.1 BrainMap database

Our analysis is motivated by coordinate data from *BrainMap* ^1^ *(Laird et al*., 2005). BrainMap is an online, freely accessible database for coordinate-based data of both functional and structural neuroimaging experiments. The data are exclusively obtained from peer-reviewed papers on whole-brain, voxel-wise studies, that are written in English language. There are three possible routes via which a paper can be to added the database. Firstly, the literature is scanned for new meta-analyses not conducted in collaboration with the BrainMap team; in these cases the meta-analysis authors are contacted and their data requested for inclusion. Secondly, they are provided by researchers who conduct meta-analysis in collaboration with the BrainMap team. Finally, they are coded after recommendation by researchers through *Scribe*, a software that is maintained by BrainMap.

Thanks to these contributions, the database has been continuously expanding since being introduced in 1998. As of April 2019, it consists of results obtained from 3,502 scientific papers on functional neuroimaging, each containing several experiments or contrasts. Recently, BrainMap also introduced a voxel-based morphometry sector (Vanasse *et al*., 2018) which, as of April 2019, contains results from 992 papers. Due to its size, BrainMap is a widely used resource for neuroimaging meta-analysis. More specifically, there are currently (April 2019) 861 peer-reviewed articles using the BrainMap and/or its CBMA software. Some recent examples include Hill *et al*. (2014), Kirby and Robinson (2017) and Hung *et al*. (2018). Throughout this paper, we assume that the database is indicative of the population of non-null neuroimaging studies; we discuss the plausibility of this assumption in Section 4.

In this work we do not consider any of the resting-state (because this sector is currently under-represented) or structural studies registered in BrainMap. Our unit of observation is a contrast, and hence our dataset consists of 16,285 observations. Each observation (contrast) consists of a list of three dimensional coordinates ***z***_*i*_, the *foci*, typically either local maxima or centers of mass of voxel clusters with significant activations. For the purposes of this work, we do not use the coordinates, and model the file drawer solely based on the total number of foci per contrast *n*_*i*_. Table 1 presents some summary statistics for the subset of the BrainMap dataset that we use (*i*.*e*. functional experiments excluding resting state), whereas Figure 2 shows the empirical distribution of the total number of foci per contrast.

**Table 1:** BrainMap database summaries.

| Database composition |  |  |  |  |  |
| --- | --- | --- | --- | --- | --- |
| Publications |  | Contrasts |  | Foci |  |
| 3,492 |  | 16,285 |  | 127,713 |  |
| Contrasts per publication |  |  |  |  |  |
| Min. | $Q_1$ | Median | Mean | $Q_3$ | Max. |
| 1 | 2 | 4 | 4.7 | 6 | 63 |
| Foci per contrast |  |  |  |  |  |
| Min. | $Q_1$ | Median | Mean | $Q_3$ | Max. |
| 1 | 2 | 5 | 7.8 | 11 | 98 |

**Figure 2:**
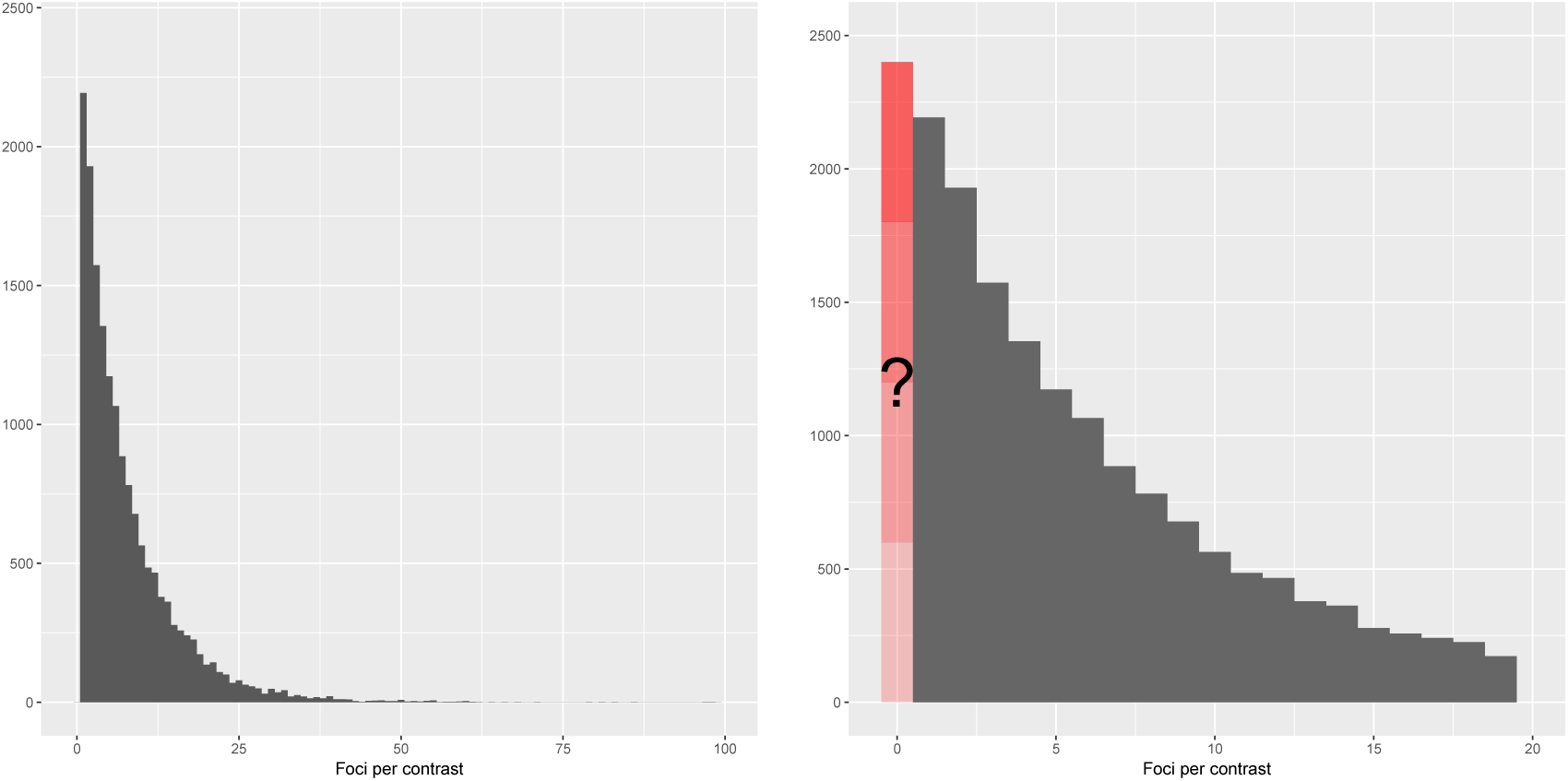
Empirical distribution of the total number of foci per contrast in the BrainMap database, *n*_*i*_. The left panel shows the full distribution, while the right panel shows a zoomed-in view of all experiments reporting 24 or fewer foci. The BrainMap database does not record incidents of null contrasts (contrasts in a paper for which *n*_*i*_ = 0).

The barplot of Figure 2 (right) identifies a fundamental aspect of this data: even though the distribution of *n*_*i*_ has most of its mass close to zero, there are no contrasts with zero foci. This is expected as by design, the contrasts from a paper that report no activations are not registered into the BrainMap database. The objective of this work is to identify the relative proportion of these contrasts compared to the ones that are registered. Some of these null contrasts may in fact be clearly reported in the papers but not registered in the BrainMap database. However, given the stigma of the negative findings, we suspect that they are rare.

### 2.2 Models

As discussed earlier, our model uses count data from the observed, reported experiments to infer on the file drawer quantity. At this point, we list the two critical assumptions: I) data 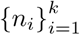, both observed and unobserved, are taken to be independent and identically distributed (i.i.d.) samples from a count distribution *N* of a given parametric form (we will relax this assumption later, to allow for inter-experiment covariates); II) the probability of publication equals zero for experiments (contrasts) with *n*_*i*_ = 0. Assumption II implies that a paper will not appear in BrainMap only if all its contrasts are negative. For a detailed discussion of the implications of assumptions I-II, see Section 4.

As each paper in the BrainMap database has multiple contrasts, potentially violating the independence assumption, we draw subsamples such that exactly one contrast from each publication is used. Specifically, we create 5 subsamples (A-E) drawing 5 different contrasts for each subsample, if possible; for publications with less than 5 contrasts we ensure that every contrast is used in at least one subsample, and then randomly select one for the remaining subsamples.

If assumptions I-II described above hold, then a suitable model for the data is a *zero-truncated* count distribution. A zero-truncated count distribution occurs when we restrict the support of a count distribution to the positive integers. For a probability mass function (pmf) *π* (*n* | ***θ***) defined on *n* = 0, 1, …, where ***θ*** is the parameter vector, the zero truncated pmf is:

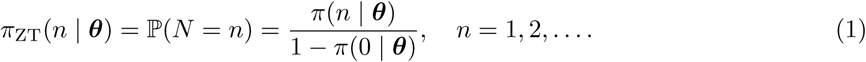

We consider three types of count distributions *π* (*n* | ***θ***): the Poisson, the Negative Binomial and the Delaporte. The Poisson is the classic distribution for counts arising from series of independent events. In particular, if the foci in a set of experiments arise from a spatial Poisson process with common intensity function, then the resulting counts will follow a Poisson distribution. Poisson models often fit count data poorly due to *over-dispersion*, that is, the observed variability of the counts is higher than what would be anticipated by a Poisson distribution. More specifically, if a spatial point process has a random intensity function, one that changes with each experiment, the distribution of counts will show this over-dispersion.

The Negative Binomial distribution is the count distribution arising from the Poisson-Gamma mixture: if the true Poisson rate differs between experiments and is distributed as a Gamma random variable, then the resulting counts will follow a Negative Binomial distribution. For the Negative Binomial distribution we use the mean-dispersion parametrisation:

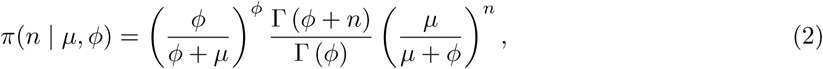

where *µ* is the mean, *ϕ* > 0 is the dispersion parameter and Γ (·) represents the gamma function; with this parametrisation the variance is 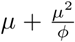. Hence, the excess of variability compared to the Poisson model is accounted for through the additional term 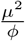.

The Delaporte distribution is obtained by modelling the foci counts *n*_*i*_ of experiment *i* as Pois(*µγ*_*i*_) random variables; the *γ*_*i*_ follows a particular shifted Gamma distribution with parameters *σ* and *ν, σ* > 0 and 0 ≤ *ν* < 1 (Rigby *et al*., 2008). The probability mass function of the Delaporte distribution can be written as:

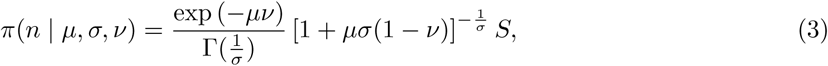

where *µ* is the mean and:

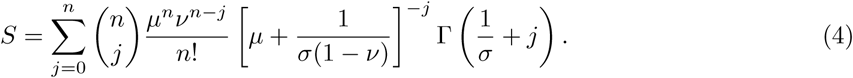

With this parametrisation the variance of the Delaporte distribution is *µ* + *µ*^2^*σ*(1 − *ν*)^2^.

Once the parameters of the truncated distribution are estimated, one can make statements about the original, untruncated distribution. One possible way to express the file drawer quantity that we are interested in is the percent prevalence of zero count contrasts *p*_z_, that is, the total number of missing experiments per 100 published. This can be estimated as:

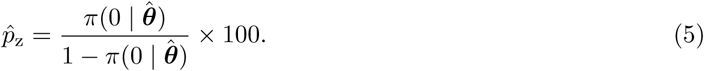

Here, 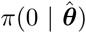 denotes the probability of observing a zero count contrast, and 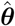 denotes the estimated parameter values from the truncated model (e.g. ***θ*** = (*µ, σ, ν*)^⊤^ for the Delaporte model).

Our statistical model is based on homogeneous data, and we can reasonably expect that differences in experiment type, sample size, etc., can introduce systematic differences between experiments. To explain as much of this nuisance variability as possible, we further model the expected number of foci per experiment as a function of its characteristics in a log-linear regression:

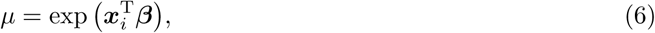

where **x**_*i*_ is the vector of covariates and ***β*** is the vector of regression coefficients. The covariates that we consider are: i) the year of publication ranging from 1985 to 2018; ii) the square root of the number of participants^2^ ranging from 1 to 395; iii) the experimental context. In each subsample, we merge all the labels of the variable context that are missing or appear less than 20 times into the ‘Other’ category. The remaining categories (that appear in at least one subsample) are: aging, disease, disease/emotion, disease/pharmacology, disease/treatment, experimental design/normal mapping, gender, language, learning, normal mapping and pharmacology. Summaries of the BrainMap subsamples A-E data for each level of context can be found in A.

Parameter estimation is done under under the *generalized additive models for location scale and shape* (GAMLSS) framework of Rigby and Stasinopoulos (2005). The fitting is done in R^3^ (R Core Team, 2015) with the *gamlss* library (Stasinopoulos and Rigby, 2007). Confidence intervals are obtained with the bootstrap. When covariates are included in the model, we use the stratified bootstrap to ensure representation of all levels of the experimental context variable. In particular, for each level of the categorical variable a bootstrap subsample is drawn using the data available for this class and subsequently these subsamples are merged to provide the bootstrap dataset. Model comparison is done using the Akaike information criterion (AIC) and the Bayesian information criterion (BIC) provided by the package.

### 2.3 Monte Carlo evaluations

We perform a simulation study to assess the quality of estimates of *p*_z_, the total number of experiments missing per 100 published, obtained by the zero-truncated Negative Binomial and Delaporte models (initial work found BrainMap counts completely incompatible with the Poisson model, and hence we did not consider it for simulation). For both approaches, synthetic data are generated as follows. First, we fix the values of the parameters, that is, *µ, ϕ* for the Negative Binomial distribution and *µ, σ, ν* for the Delaporte distribution. We then generate *k*\**/*(1 − *π*(0|***θ***)) samples from the untruncated distributions, where *k*\* is chosen such that the expected number of non-zero counts is *k*. We remove the zero-count instances from the simulated data and the corresponding zero-truncated model is fit to the remaining observations. Finally, we estimate the probability of observing a zero count experiment based on our parameter estimates.

We set our simulation parameter values to cover typical values found in BrainMap (see C, Table 8). For the Negative Binomial distribution we consider values 4 and 8 for the mean and values 0.4, 0.8, 1.0 and 1.5 for the dispersion, for a total of 8 parameter settings. For the Delaporte distribution, we set *µ* to 4 and 8, *σ* to 0.5, 0.9 and 1.2, and *ν* to 0.02, 0.06 and 0.1 (18 parameter settings). The expected number of observed experiments is set to *k* = 200, 500, 1,000 and 2,000. For each combination of (*k, µ, ϕ*) and (*k, µ, σ, ν*) of the Negative Binomial and Delaporte models, respectively, we generate 1,000 datasets from the corresponding model, for each parameter setting, and record the estimated value of *p*_z_ for each fitted dataset.

### 2.4 HCP real data evaluations

As an evaluation of our methods on realistic data for which the exact number of missing contrasts is known, we generate synthetic meta-analysis datasets using the Human Connectome Project task fMRI data. We start with a selection of 80 unrelated subjects and retrieve data for all 86 tasks considered in the experiment. For each task, we randomly split the 80 subjects into 8 groups of 10 subjects. Hence, we obtain a total of 86 × 8 = 688 synthetic fMRI experiments. For each experiment, we perform a one-sample group analysis, using ordinary least squares in FSL^4^, and recording 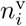, the total number of surviving peaks after random field theory thresholding at the voxel level, 1% familywise error rate (FWE), where *i* = 1, …, 688. We also record the total number of peaks (one peak per cluster) after random field theory thresholding at the cluster level, cluster forming threshold of uncorrected P=0.00001 & 1% FWE, 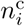. These rather stringent significance levels were needed to induce sufficient numbers of results with no activations. We then discard the zero-count instances from 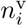 and 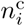, and subsequently analyse the two truncated samples in two separate analyses, using the zero-truncated Negative Binomial and Delaporte models. Finally, the estimated number of missing experiments is compared to the actual number of discarded contrasts. Note that we repeat the procedure described above 6 times, each time using different random splits of the 80 subjects (HCP splits 1-6).

## 3 Results

### 3.1 Simulation results

The percent relative bias of the estimates of 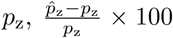, and its bootstrap standard error for the zero-truncated Negative Binomial and Delaporte models are shown in Table 2 and Table 3, respectively. The results indicate that, when the model is correctly specified, both approaches perform adequately. In particular, in Table 2 we see that the bias of 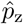 is small, never exceeding 8% when the sample size is comparable to the sample size of the BrainMap database (*k* = 3, 492) and the mean number of foci is similar to the average foci count found in BrainMap (≈ 9). The bootstrap standard error estimates produced by the Negative Binomial model are also accurate with relative bias below 5% in most scenarios with more than 500 contrasts, while Delaporte tends to underestimate standard errors but never more than −15% (see Table 3).

**Table 2:** Percent relative bias for estimation of *p*_z_, the zero-count experiment rate as a percentage of observed experiments, for Negative Binomial and Delaporte models as obtained from 1,000 simulated datasets. Parameter *µ* is the expected number of foci per experiment, *ϕ, σ* and *ν* are additional scale and shape parameters. Negative Binomial performs well and, while Delaporte often underestimated *p*_z_, with at least 1,000 contrasts it always has bias less than 10% (positive bias over-estimates the file drawer problem).

| Negative Binomial |  |
| --- | --- |
| Parameter values | % relative bias of $\hat{p}_z$ |

| $\mu$ | $\phi$ | $p_z$ | $\mathbb{E}[k] = 200$ | 500 | 1000 | 2000 |
| --- | --- | --- | --- | --- | --- | --- |
| 4 | 0.4 | 62.1 | 8.76 | 2.85 | 1.40 | 0.80 |
| 4 | 0.8 | 31.3 | 2.72 | 1.97 | -0.78 | 0.32 |
| 4 | 1.0 | 25.0 | 1.70 | 1.21 | 0.72 | 0.16 |
| 4 | 1.5 | 16.6 | 2.85 | 1.66 | 0.71 | 0.13 |
| 8 | 0.4 | 42.0 | 7.07 | 3.17 | 0.70 | -0.22 |
| 8 | 0.8 | 17.2 | 4.35 | 1.57 | 0.49 | 0.21 |
| 8 | 1.0 | 12.5 | 3.32 | 0.84 | 0.63 | 0.04 |
| 8 | 1.5 | 6.7 | 2.53 | 0.90 | 0.69 | 0.21 |

**Delaporte**
| Parameter values | | | | % relative bias of $\hat{p}_z$ | | | |
| --- | --- | --- | --- | --- | --- | --- | --- |
| $\mu$ | $\sigma$ | $\nu$ | $p_z$ | $\mathbb{E}[k] = 200$ | 500 | 1000 | 2000 |
| 4 | 0.5 | 0.02 | 11.8 | -12.65 | -10.66 | -9.69 | -8.02 |
| 4 | 0.9 | 0.02 | 20.8 | -17.47 | -12.35 | -10.28 | -10.10 |
| 4 | 1.2 | 0.02 | 27.6 | -20.46 | -18.59 | -16.64 | -13.83 |
| 4 | 0.5 | 0.06 | 10.5 | -6.13 | -5.58 | -4.77 | -3.73 |
| 4 | 0.9 | 0.06 | 18.0 | -4.08 | -4.18 | -1.32 | 0.15 |
| 4 | 1.2 | 0.06 | 23.4 | -5.07 | -3.27 | -1.09 | 0.01 |
| 4 | 0.5 | 0.10 | 9.3 | -4.53 | -3.32 | -2.65 | -1.90 |
| 4 | 0.9 | 0.10 | 15.6 | 1.07 | 3.86 | 1.91 | 1.80 |
| 4 | 1.2 | 0.10 | 20.0 | 4.46 | 3.32 | 3.89 | 4.07 |
| 8 | 0.5 | 0.02 | 3.6 | -13.77 | -10.02 | -7.75 | -5.91 |
| 8 | 0.9 | 0.02 | 9.2 | -11.99 | -9.00 | -7.46 | -5.04 |
| 8 | 1.2 | 0.02 | 13.8 | -14.07 | -11.22 | -9.48 | -8.12 |
| 8 | 0.5 | 0.06 | 2.8 | -3.04 | -3.18 | -2.12 | -1.89 |
| 8 | 0.9 | 0.06 | 6.8 | 1.41 | 4.13 | 2.74 | 2.47 |
| 8 | 1.2 | 0.06 | 10.0 | 10.49 | 7.52 | 7.91 | 5.19 |
| 8 | 0.5 | 0.10 | 2.2 | 0.93 | 1.36 | 0.82 | 0.74 |
| 8 | 0.9 | 0.10 | 5.0 | 8.09 | 5.88 | 5.78 | 4.94 |
| 8 | 1.2 | 0.10 | 7.3 | 17.91 | 10.68 | 7.77 | 4.76 |

**Table 3:**
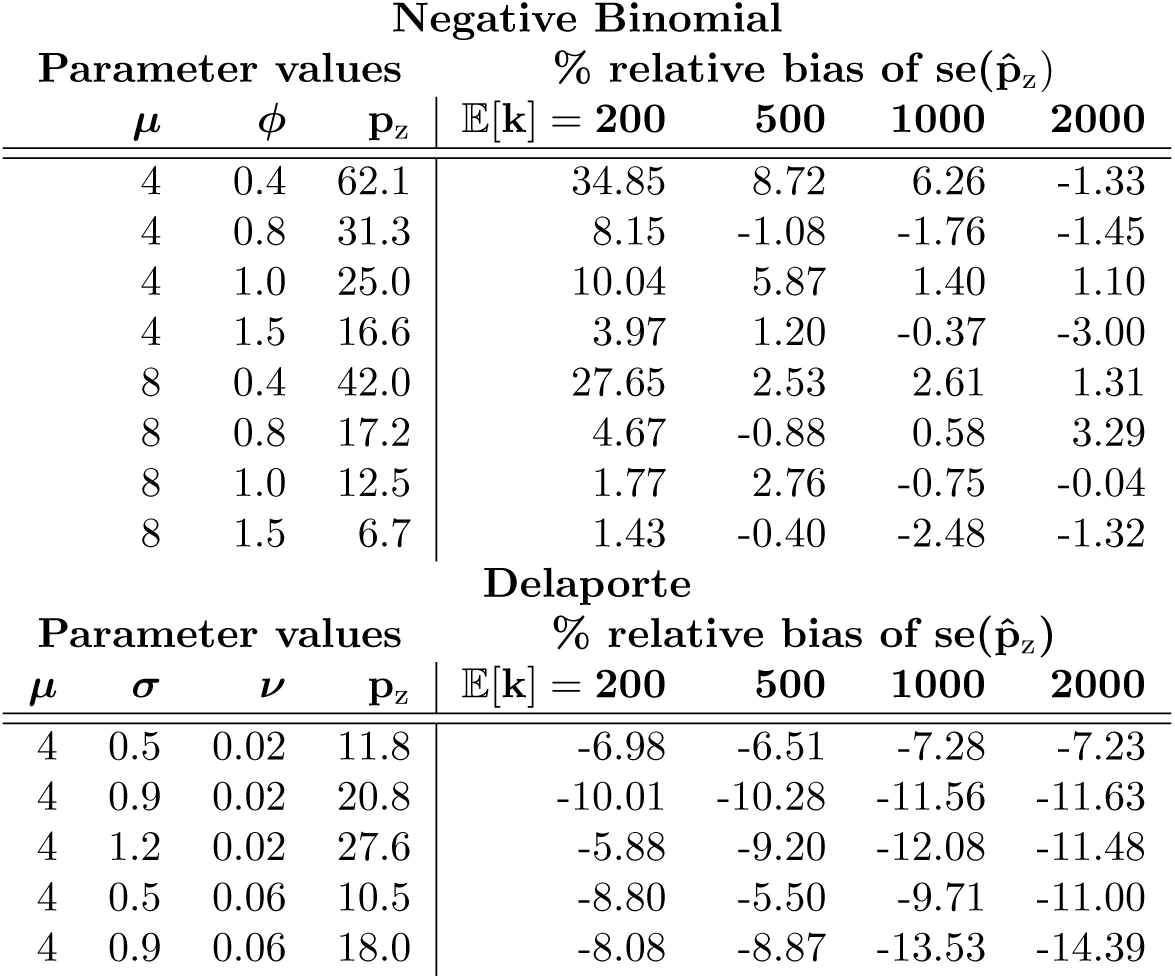
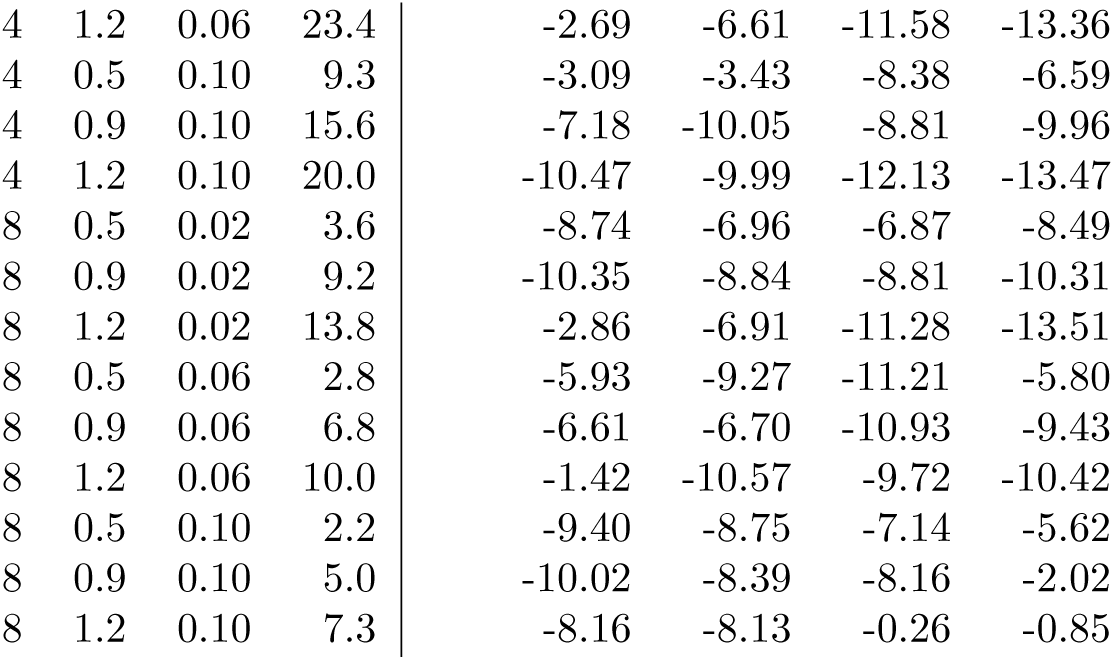
Percent relative bias of bootstrap standard error of 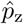, missing experiment rate as a percentage of observed experiments, for Negative Binomial and Delaporte models as obtained from 1,000 simulated datasets. Parameter *μ* is the expected number of foci per experiment and *ϕ, σ* and *ν* are additional scale and shape parameters. For a sample of at least 1,000 contrasts, Negative Binomial standard errors are usually less than 3% in absolute value; while Delaporte has worse bias, it is never less than −15% (negative standard error bias leads to over-confident inferences).

### 3.2 HCP synthetic data results

Results of the analysis of the HCP synthetic datasets using the zero-truncated Negative Binomial and Delaporte models are summarised in Figure 3 and Table 4. In Figure 3 we plot the empirical count distributions and the fitted probability mass functions for the 12 datasets considered. For datasets obtained with voxelwise thesholding of the image data, we see that the Delaporte distribution provides a better fit compared to the Negative Binomial qualitatively, by AIC in all 6 datasets, and by BIC in five out of six datasets (Table 4). For clusterwise thresholding, there are fewer peaks in general and their distribution is less variable compared to voxelwise thresholding. Both distributions achieve a similar fit. Here, AIC supports the Negative Binomial model in 4 out of 6 datasets and BIC in 5 out of 6 datasets.

**Table 4:**
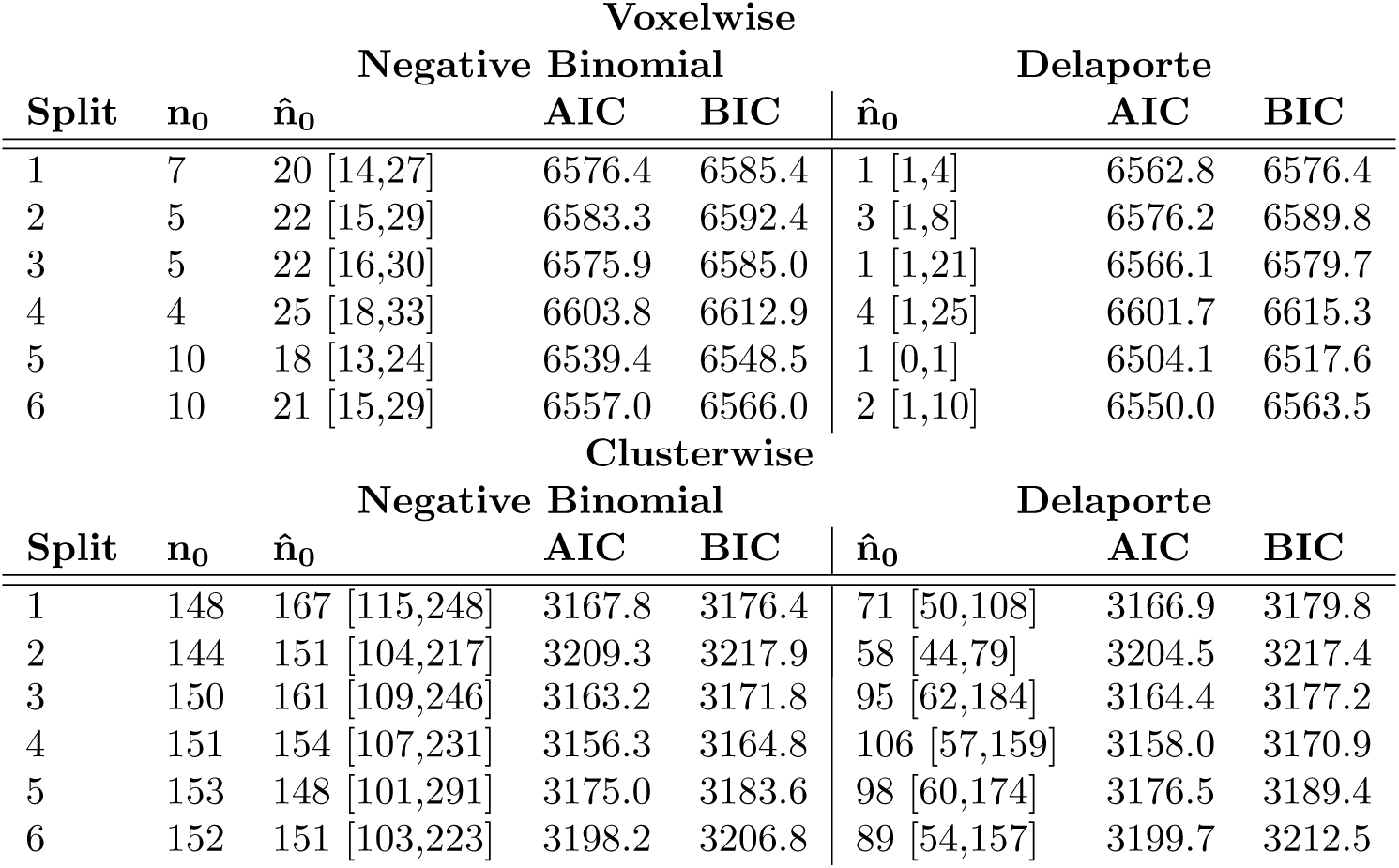
Evaluation of the zero-truncated modelling approach using synthetic data obtained from the HCP project, using voxelwise (top) and clusterwise (bottom) inference. The true number of missing contrasts (*n*_0_) for each one of the 12 datasets (6 for voxelwise thesholding and 6 for clusterwise thresholding) is shown in the second column. For each of the Negative Binomial and Delaporte methods, the estimated missing contrast count 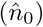, 95% bootstrap confidence interval for *n*_0_, AIC score and BIC score are shown (smaller AIC and BIC are better).

**Figure 3:**
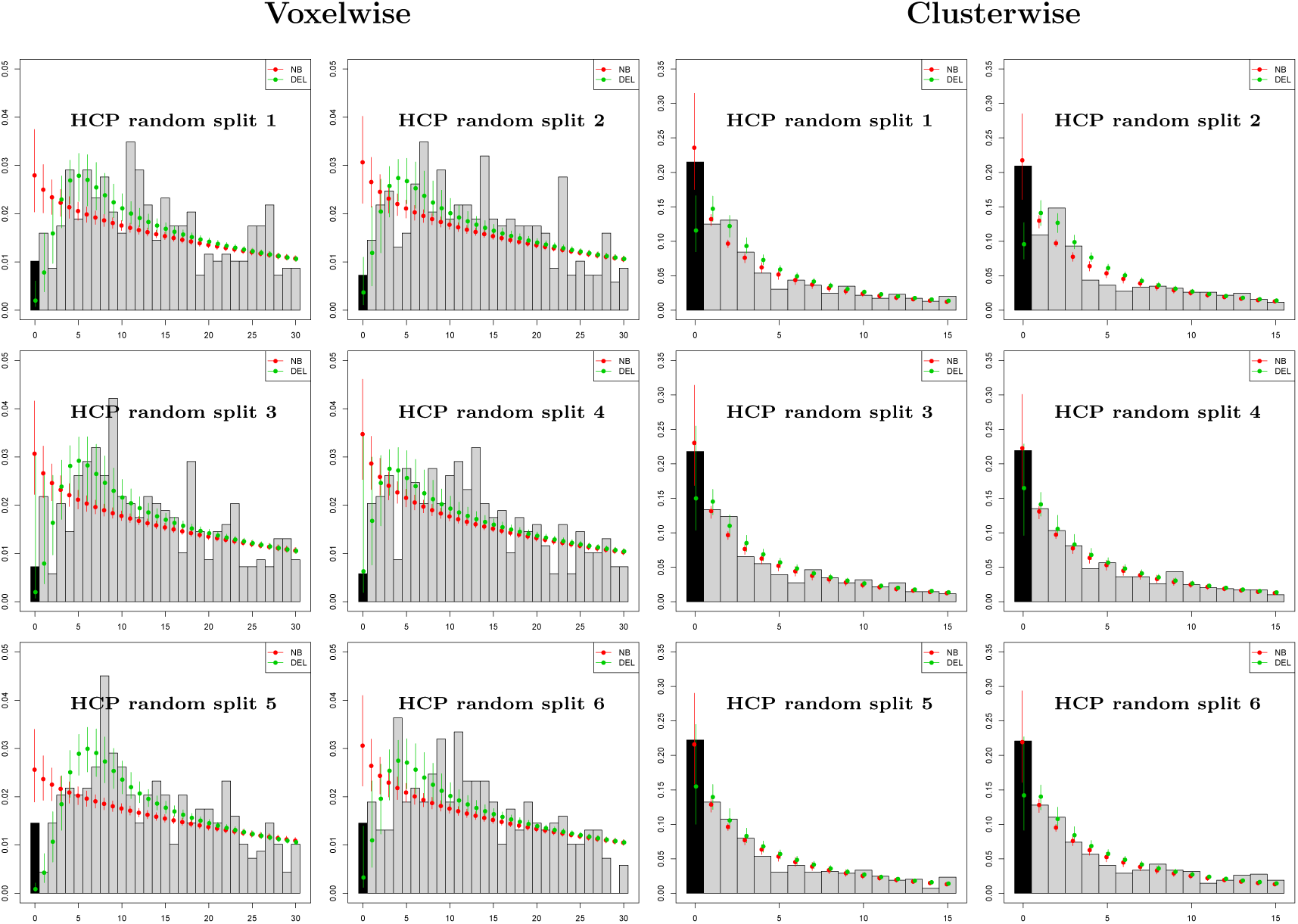
Evaluation with HCP data with 688 contrasts of sample size 10, comparing accuracy of Negative Binomial (NB) and Delaporte (DEL) distributions for the prediction of the number of contrasts with no significant results (zero foci) based on only significant results (one or more foci). Left panel shows results for voxelwise inference, right for clusterwise inference, both using *P*_FWE_=0.01 to increase frequency of zero foci. For clusterwise datasets, the Negative Binomial confidence intervals always include the observed zero count, while Delaporte ofter underestimates the count. For voxelwise analysis, the Negative Binomial over-estimates the zero frequency substantially, while Delaporte’s intervals include the actual zero frequency in 3 out of 5 splits.

Table 4 reports the true number of missing contrasts *n*_0_, along with point estimates 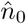 and the 95% bootstrap intervals obtained by the zero-truncated models. For voxelwise data, the Negative Binomial model overestimates the total number of missing experiments in all 6 datasets as a consequence of the poor fit to the non-zero counts, while the Delaporte model bootstrap intervals include the true value of *n*_0_ in 5 out of 6 datasets, greatly underestimating *n*_0_ in one dataset. For clusterwise counts, the point estimates obtained by the zero-truncated Negative Binomial model are very close to the true values. Notably, *n*_0_ is included within the bootstrap intervals for all 6 datasets. The Delaporte model underestimates the values of *n*_0_ in all 6 datasets, but the bootstrap intervals include *n*_0_ for 4 out of 6 datasets.

Overall, we find that the zero-truncated modeling approach generally provides good estimates of *n*_0_, with the Negative Binomial sometimes overestimating and the Delaporte sometimes underestimating *n*_0_. A conservative approach, therefore, favors the Delaporte model.

### 3.3 Application to the BrainMap data

We found the Poisson distribution to be completely incompatible with the BrainMap count data (B, Figure 8), and we do not consider it further. We start by fitting the Negative Binomial and Delaporte zero-truncated models without any covariates. The estimates of the scalar parameters obtained for both models are shown in Appendix C. Figure 4 shows the emprical and fitted probability mass functions for the 5 subsamples. We see that both distributions provide a good fit for the BrainMap data. The Negative Binomial model is preferred based on AIC in 4 out of 5 subsamples, and based on BIC in 5 out of 5 subsamples (Table 6), buth with little difference in both criteria. The estimated prevalence of missing contrasts, along with 95% bootstrap intervals are shown in Table 5. Note that while there is considerable variation in the estimates over the two models, the confidence intervals from all subsamples do not include zero, thus suggesting a file drawer effect.

**Table 5:** BrainMap data analysis results. The table presents the estimated prevelance of file drawer experiments along with 95% bootstrap confidence intervals, as obtained by fitting the zero-truncated Negative Binomial and Delaporte models to BrainMap subsmaples A-E. No covariates are considered.

| Subsample | Negative Binomial |  | Delaporte |  |
| --- | --- | --- | --- | --- |
| | $\hat{p}_z$ | 95% interval | $\hat{p}_z$ | 95% interval |
| A | 11.80 | [10.22,13.59] | 10.52 | [6.42,13.04] |
| B | 11.14 | [9.63,12.79] | 7.30 | [5.29,11.44] |
| C | 11.46 | [9.95,13.31] | 8.30 | [5.95,12.04] |
| D | 11.16 | [9.64,12.88] | 6.83 | [5.03,10.97] |
| E | 11.40 | [9.89,13.14] | 8.11 | [5.44,12.30] |

**Table 6:** AIC/BIC model comparison results for the BrainMap data. We fit the zero-truncated Negative Binomial and Delaporte models, with and without the covariates, to BrainMap subsamples A-E. Every split indicates evidence for better fit with the Negative Binomial model (smaller AIC/BIC indicates better fitting model). For regression models, the sample size was smaller due to missing values. Hence, the criteria cannot be used to compare the models without and with the covariates.

| Model comparison: no covariates |  |  |  |  |
| --- | --- | --- | --- | --- |
| Subsample | Negative Binomial |  | Delaporte |  |
|  | AIC | BIC | AIC | BIC |
| A | 21837.87 | 21850.18 | 21839.87 | 21858.34 |
| B | 21792.04 | 21804.36 | 21792.61 | 21811.09 |
| C | 21899.71 | 21912.03 | 21901.18 | 21919.65 |
| D | 21862.23 | 21874.54 | 21861.83 | 21880.30 |
| E | 21683.47 | 21695.79 | 21684.94 | 21703.42 |

| Model comparison: regression |  |  |  |  |
| --- | --- | --- | --- | --- |
| Subsample | Negative Binomial |  | Delaporte |  |
|  | AIC | BIC | AIC | BIC |
| A | 21778.73 | 21871.10 | 21780.45 | 21878.98 |
| B | 21735.59 | 21827.96 | 21733.23 | 21831.76 |
| C | 21850.23 | 21936.44 | 21850.29 | 21942.66 |
| D | 21816.27 | 21902.49 | 21813.07 | 21905.45 |
| E | 21628.24 | 21714.45 | 21628.58 | 21720.95 |

**Figure 4:**
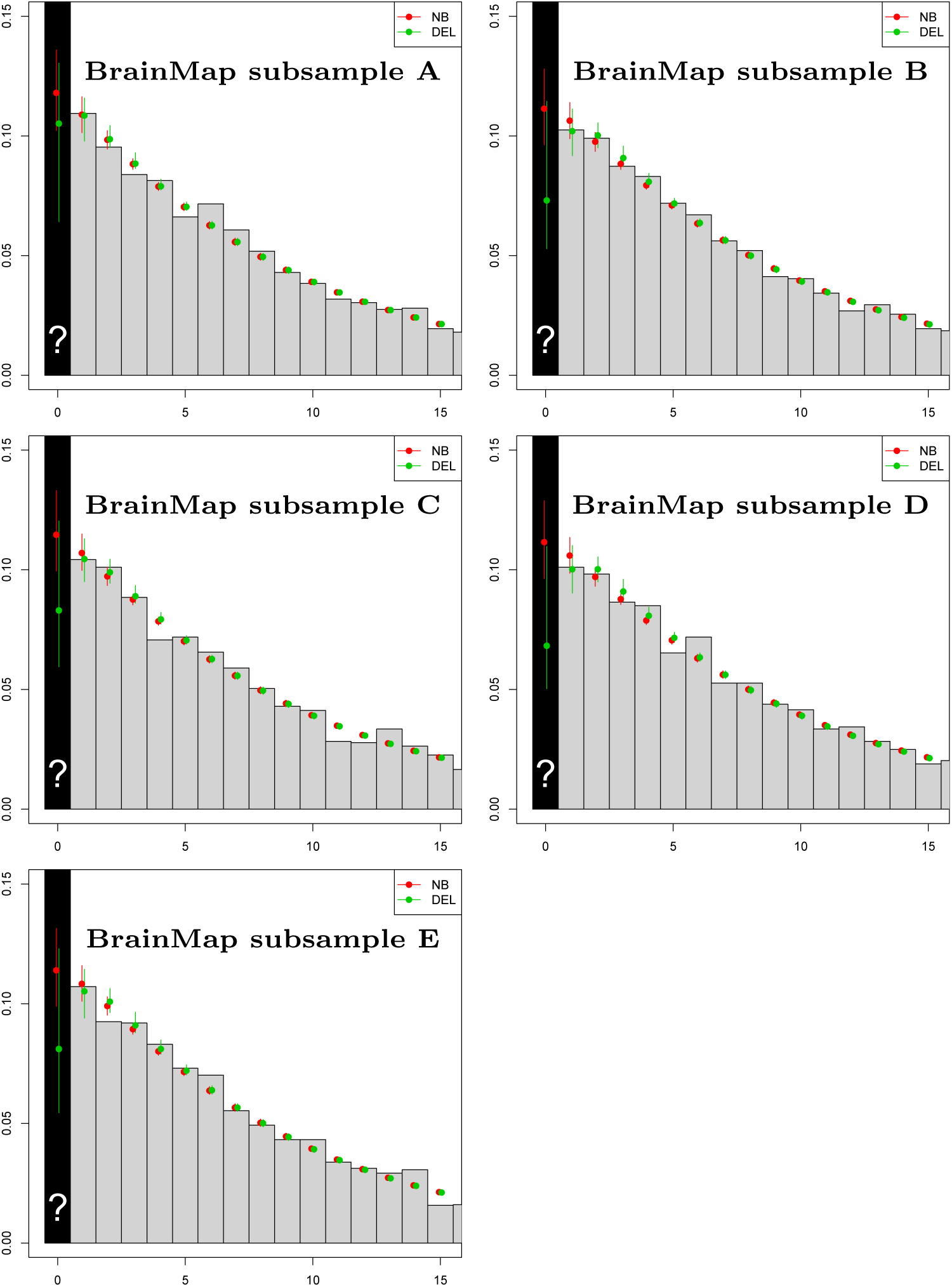
BrainMap results for 5 random samples using the Negative Binomial and Delaporte models and no covariates. Plots show observed count data (gray bars) with fit of full (non-truncated) distribution based on zero-truncated data, including the estimate of *p*_0_ (over black bar).

For both Negative Binomial and Delaporte models, and all subsamples A-E, the model with covariates is preferred over the simple model (without the covariates) in terms of AIC but not in terms of BIC (Table 6). This is expected since BIC penalizes model complexity more heavily. Covariates essentially have no effect on the estimated prevalence of missing contrasts. As can be seen in Figure 5, the estimated prevalence of zero count contrasts is a slowly decreasing function of both the square root number of participants and the year of publication. For the former, the trend is expected and one possible explanation is that bigger samples result into greater power, and therefore more foci and thus less of a file drawer problem. However, for publication year, decreasing publication bias is welcomed but we could have just as well expected that the increased use of multiple testing in later years would have reduced foci counts and *increased* the file drawer problem. We further see that the estimated percent prevalence of zero-count contrasts is similar for all levels of the categorical variable context, with the exception of experiments studying gender effects (Figure 6). Finally, when including the covariates, the Negative Binomial is preferred over the Delaporte in 3 out of 5 subsamples in terms of the AIC, and in 4 out of 5 subsamples in terms of the BIC (Table 6).

**Figure 5:**
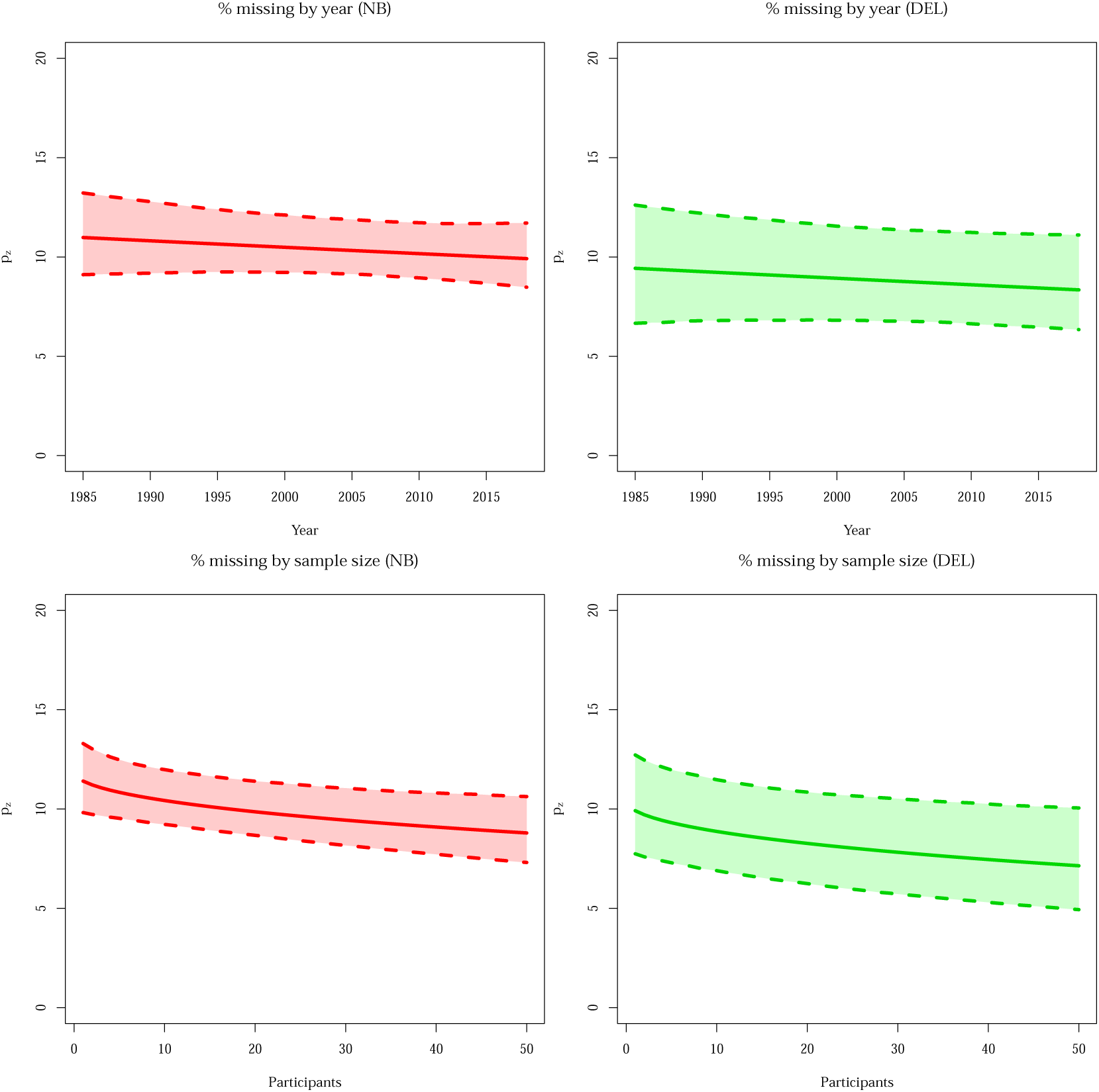
Predicted *p*_z_, missing experiment rate per 100 published experiments, as a function of year of publication (top) and the square root of sample size (bottom), with pointwise 95% bootstrap confidence intervals. There is not much variation in the estimate of the percentage missing, but in both cases a negative slope is observed, as might be expected with improving research practices over time and greater power with increased sample size. All panels refer to the second BrainMap random sample (subsample B).

**Figure 6:**
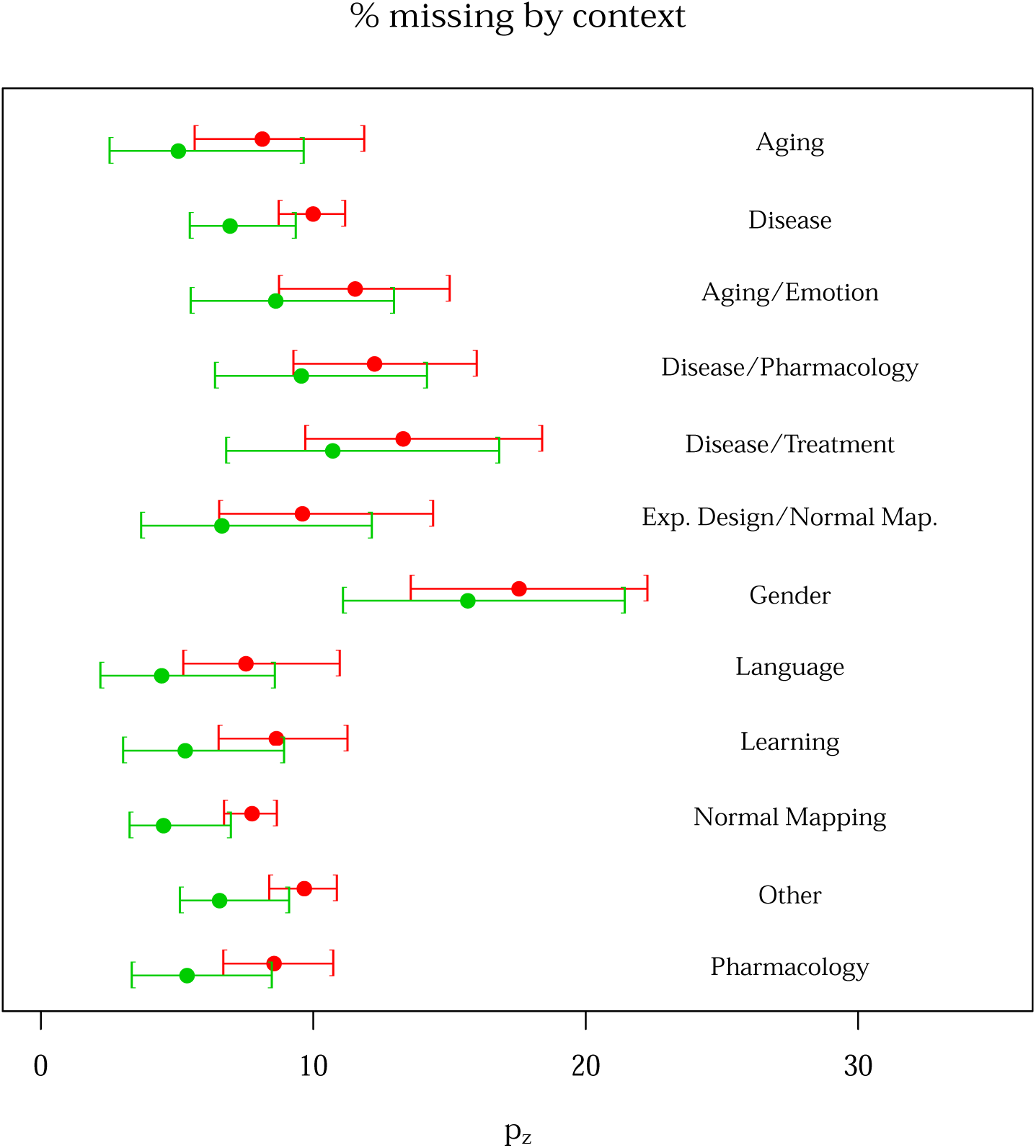
Contrasts missing per 100 published as a function of experiment context, with 95% bootstrap confidence intervals. Note that we have fixed the year and square root sample size covariates to their median values. The plot is derived from the second BrainMap random sample (subsample B). The red and green lines represent the Negative Binomial and Delaporte distributions, respectively.

## 4 Discussion

### 4.1 Summary of findings & implications for CBMA

In this paper, we have attempted to estimate the prevalence of experiments missing from a CBMA due to reporting non-significant results. Our method uses intrinsic statistical characteristics of the non-zero count data to infer the relative frequency of zero counts. This is achieved by estimating the parameters of a zero-truncated model, either Negative Binomial or Delaporte, which are subsequently used to predict the prevalence *p*_0_ of zero-count experiments in the original, untruncated distribution, and re-expressing this as *p*_z_, the rate of missing contrasts per 100 published.

Our approach further relies on assumptions I and II described in Section 2.2. Assumption I implies that there is independence between contrasts. However, as one publication can have several contrasts, this assumption is tenuous despite it being a standard assumption for most CBMA methods. To ensure the independence assumption is valid, we subsample the data so that only one randomly selected contrast per publication is used. Assumption II defines our censoring mechanism, such that only experiments with at least one significant activation can be published. The assumption that non-significant research findings are suppressed from the literature has been adopted by authors in classical meta-analysis (Eberly and Casella, 1999, among others). One possible way in which this assumption can be violated could be due to data repeatedly analysed under different pipelines (e.g. by using a different cluster extent each time) until they provide significant results. However, we believe that are unlikely to resort to this approach because studies typically involve multiple contrasts. Hence, even if some of them are negative, the authors can focus on remaining, significant contrasts, in their publication.

A series of simulations studies suggest that the zero-truncated modelling approach provides valid estimates of *p*_z_. A critical limitation of our HCP evaluation is the repeated measures structure, where 86 contrasts come from each subject. Such dependence generally does not induce bias in the mean estimates, but can corrupt standard errors and is a violation of the bootstrap’s independence assumption. However, as the bootstrap intervals generally captured the true censoring rate, it seems we were not adversely affected by this violation. It should be noted, moreover, that the properties of our estimators degrade as the total number of observed experiments decreases and therefore our methods are likely not suitable for individual meta-analyses unless hundreds of experiments are available.

The analysis of BrainMap data suggests that the estimated prevalence of null contrasts slightly varies depending on the characteristics of an experiment, but generally consists of at least 6 missing experiments for 100 published, and this estimate of 6 is significantly greater than zero. In other words, for a randomly selected CBMA consisting of *J* contrasts, we expect that 6*J*/100 null contrasts are missing due to the file drawer. Note that this interpretation concerns the aggregate statistical practice reflected in BrainMap, i.e. it is totally agnostic to the statistical procedures used to generate the results in the database. The counts we model could have been found with liberal *P* < 0.001 uncorrected inferences or stringent *P* < 0.05 FWE procedures. However, if the neuroimaging community *never* used multiple testing corrections, then every experiment should report many peaks, and we should estimate virtually no missing contrasts.

The results suggest that the population sampled by the BrainMap database has a non-zero file drawer effect. Whether this conclusion can be extended to all neuroimaging experiments depends on the representativeness of the database. As noted above, the BrainMap staff are continually adding studies and capture the content of newly published meta-analyses. Hence, the most notable bias could be topicality and novelty effects that drive researchers to create meta-analyses. Another potential source of bias could be due to studies which BrainMap does not have access to, such as ones that are never published due insufficient time to submit a paper or staff leaving. But we do not see these particular effects as driving the file drawer effect up or down in particular, and so is not so much of a concern.

Our findings provide evidence for the existence of publication bias in CBMA. The presence of missing experiments does not invalidate existing CBMA studies, but complements the picture seen when conducting a literature review. Considering the missing contrasts would affect the conclusions drawn from any of the existing CBMA approaches, both model-based (such as Kang *et al*. (2014) or Montagna *et al*. (2018)) and kernel-based (such as ALE (Eickhoff *et al*., 2012) or MKDA Wager *et al*. (2007)). More specifically, the inclusion of null contrasts would decrease the estimated effect size at each voxel.

### 4.2 Future work

There are a few limitations to our work. Even though we posit that the majority of the missing contrasts are never described in publications or not published at all, we cannot rule out the contribution from contrasts that have actually been reported in the original publications and simply not encoded in BrainMap. Therefore, it is worth considering an extensive literature review in order to investigate how often such null results are mentioned in papers. This information can be then used to approximate the fraction of contrasts that are never published. Ideally, our unit of inference would be a publication rather than a contrast. However, linking our contrast-level inferences to studies requires assumptions about dependence of contrasts within a study and the distribution of the number of contrasts examined per study. We can assert that the more contrasts examined per investigation, the more likely 1 or more null contrasts should arise; and that the risk of null contrasts is inversely related to foci-per-contrast of non-null contrasts. Using these facts, there might be ways in which one could use our findings to estimate the rate of missing studies.

The evaluation of our methods using the HCP data could be also extended. One option would be to implement a different analysis pipeline in each one of the synthetic experiments in order to reflect the heterogeneity observed in the BrainMap data. Another option would be to investigate the robustness of the results to the choice of the sample size of each synthetic experiment. Nonetheless, for larger sample sizes, this would require a larger selection of HCP subjects in order to ensure that there total number of synthetic experiments is sufficient for our approach. The analysis conducted in this paper is based on data retrieved from a single database. As a consequence, results are not robust to possible biases in the way publications are included in this particular database. A more thorough analysis would require consideration of other databases (e.g. Neurosynth.org^5^ (Yarkoni *et al*., 2011), though note Neurosynth does not report foci per contrast but per paper).

Finally, one may argue that our censoring mechanism is rather simplistic, and does not reflect the complexity of current (and potentially) poor scientific practice. As discussed earlier, we have not allowed for the possibility of ‘vibration effects’, that is, changing the analysis pipeline (e.g., random vs fixed effects, linear vs. nonlinear registration) to finally obtain some significant activations. This would be an instance of initially-censored (zero-count) data being ‘promoted’ to a non-zero count through some means, see Figure 7 for a graphical representation. Such models can be fit under the Bayesian paradigm and we will consider them in our future work.

**Figure 7:**
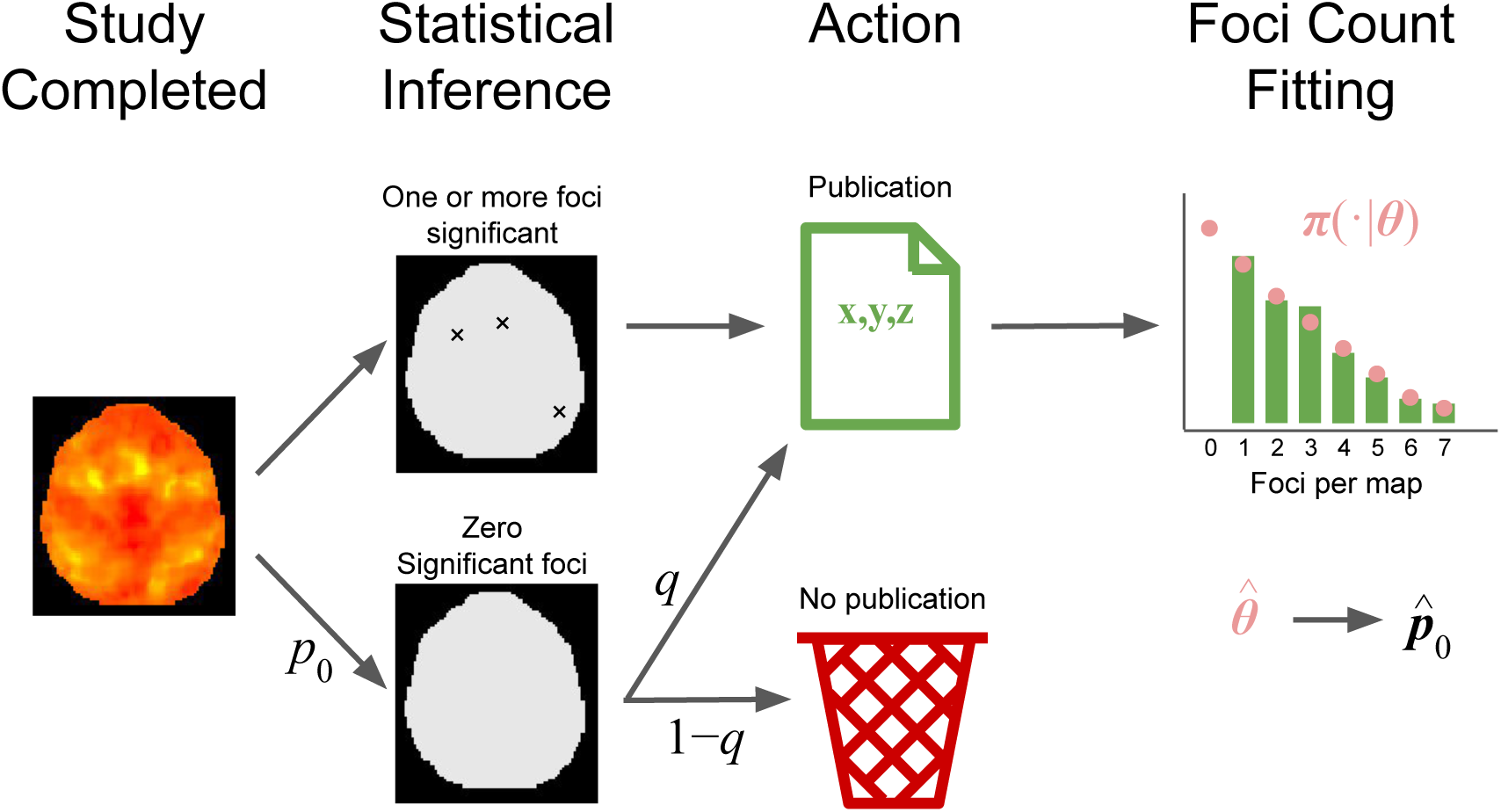
A zero-upgrade model.

Our simulation studies have shown that the properties of our prevalence estimator are poor when the total number of non-zero experiments available is low. This fact implies that the estimator cannot be used to infer the number of missing experiments from a single CBMA. Hence, a potential direction for future work would be to construct estimators that are more robust when the sample size is small. Finally, given results in this paper, there are potential benefits in extending existing CBMA methodologies to account for the file drawer. Many authors have suggested possible solutions for this problem in classic meta-analysis, using, for example *funnel plots* (Egger *et al*., 1997; Duval and Tweedie, 2000), *weight functions* (Larose and Dey, 1998; Copas and Jackson, 2004) or *sensitivity analyses* (Copas and Shi, 2000). For a recent survey of such methods, see Jin *et al*. (2015). It is therefore conceivable to adapt these methods for application in CBMA. Acar *et al*. (2018) develop a robustness check for the activation likelihood estimation algorithm (Eickhoff *et al*., 2009), which is based on the *fail-safe N* (Rosenthal, 1979; Iyengar and Greenhouse, 1988) that is, the minimum number of unpublished studies required to overturn the outcome of meta-analysis. However, it is essential that such checks are developed for other widely-used CBMA methods.

## A BrainMap summaries for context

In this section we provide summaries of the data on the 5 BrainMap subsamples A-E, for the different levels of the categorical variable experiment context. In particular, Table 7 presents the total number of contrasts per level, the average sample size per contrast, and the average number of foci per contrast. Note that in subsamples C, D and E there were less than 20 contrasts with label ‘Gender’; hence, we incorporate those in the ‘Other’ category.

**Table 7:** Data summaries for the different levels of the categorical variable experiment context.

| Contrasts per level |  |  |  |  |  |
| --- | --- | --- | --- | --- | --- |
| Experiment context | BrainMap subsample |  |  |  |  |
|  | A | B | C | D | E |
| Aging | 20 | 27 | 22 | 23 | 21 |
| Disease | 574 | 595 | 593 | 590 | 585 |
| Disease, Emotion | 34 | 32 | 31 | 33 | 31 |
| Disease, Pharmacology | 48 | 49 | 42 | 47 | 38 |
| Disease, Treatment | 33 | 30 | 31 | 33 | 28 |
| Experimental Design, Normal Mapping | 30 | 30 | 32 | 33 | 26 |
| Gender | 22 | 23 | - | - | - |
| Language | 29 | 32 | 29 | 27 | 31 |
| Learning | 26 | 31 | 25 | 27 | 28 |
| Normal Mapping | 2074 | 2043 | 2063 | 2062 | 2081 |
| Other | 539 | 542 | 563 | 555 | 566 |
| Pharmacology | 63 | 58 | 61 | 62 | 57 |
| Average contrast sample size |  |  |  |  |  |
| Experiment context | BrainMap subsample |  |  |  |  |
|  | A | B | C | D | E |
| Aging | 16.4 | 12.4 | 12.0 | 11.7 | 12.2 |
| Disease | 14.4 | 15.0 | 14.5 | 15.1 | 15.0 |
| Disease, Emotion | 15.7 | 16.1 | 16.2 | 16.0 | 16.0 |
| Disease, Pharmacology | 12.3 | 12.5 | 12.8 | 12.8 | 12.1 |
| Disease, Treatment | 11.4 | 12.9 | 12.4 | 11.9 | 11.3 |
| Experimental Design, Normal Mapping | 19.2 | 19.6 | 21.7 | 20.9 | 20.5 |
| Gender | 13.2 | 12.2 | - | - | - |
| Language | 11.6 | 11.8 | 11.6 | 11.9 | 10.7 |
| Learning | 10.4 | 11.6 | 10.2 | 10.5 | 11.8 |
| Normal Mapping | 13.9 | 13.7 | 13.8 | 13.7 | 13.9 |
| Other | 16.9 | 17.1 | 16.8 | 16.9 | 16.9 |
| Pharmacology | 13.0 | 13.4 | 12.5 | 13.2 | 12.6 |
| Average foci per contrast |  |  |  |  |  |
| Experiment context | BrainMap subsample |  |  |  |  |
|  | A | B | C | D | E |
| Aging | 10.4 | 9.1 | 8.5 | 9.2 | 6.0 |
| Disease | 7.2 | 7.5 | 7.6 | 7.5 | 7.3 |
| Disease, Emotion | 6.6 | 6.6 | 6.5 | 7.8 | 6.0 |
| Disease, Pharmacology | 5.6 | 6.0 | 6.0 | 6.6 | 5.2 |
| Disease, Treatment | 8.2 | 5.6 | 6.5 | 6.7 | 7.3 |
| Experimental Design, Normal Mapping | 7.6 | 8.0 | 7.2 | 5.9 | 8.0 |
| Gender | 5.8 | 4.0 | - | - | - |
| Language | 9.1 | 9.9 | 8.1 | 7.1 | 7.9 |
| Learning | 7.6 | 8.5 | 7.6 | 7.0 | 9.6 |
| Normal Mapping | 9.7 | 9.7 | 9.9 | 9.7 | 9.5 |
| Other | 8.1 | 7.9 | 7.8 | 8.2 | 7.9 |
| Pharmacology | 8.4 | 8.8 | 9.5 | 8.4 | 7.7 |

## B Zero-truncated Poisson analysis of the BrainMap dataset

In this section, we present results of the analysis of BrainMap subsamples A-E using the zero-truncated Poisson model. The empirical and fitted Poisson probability mass functions are shown in Figure 8. It is evident that the zero-truncated Poisson model provides a poor fit to the BrainMap data. The finding is confirmed by the AIC and BIC criteria. The AIC is 35513.5, 34886.8, 35595.9, 35456.7 and 34642.1 for subsamples A-E, respectively. The BIC is 35519.7, 34893.0, 35602.1, 35462.8 and 34648.3 for subsamples A-E, respectively. These values are much higher than the corresponding values obtained by fitting both the Negative Binomial and Delaporte models (see Table 6). The estimated prevalence of file drawer experiments is estimated as almost zero in all subsamples (Figure 8, final plot). However, these estimates should not be trusted considering the poor fit provided by the zero-truncated Poisson model.

**Figure 8:**
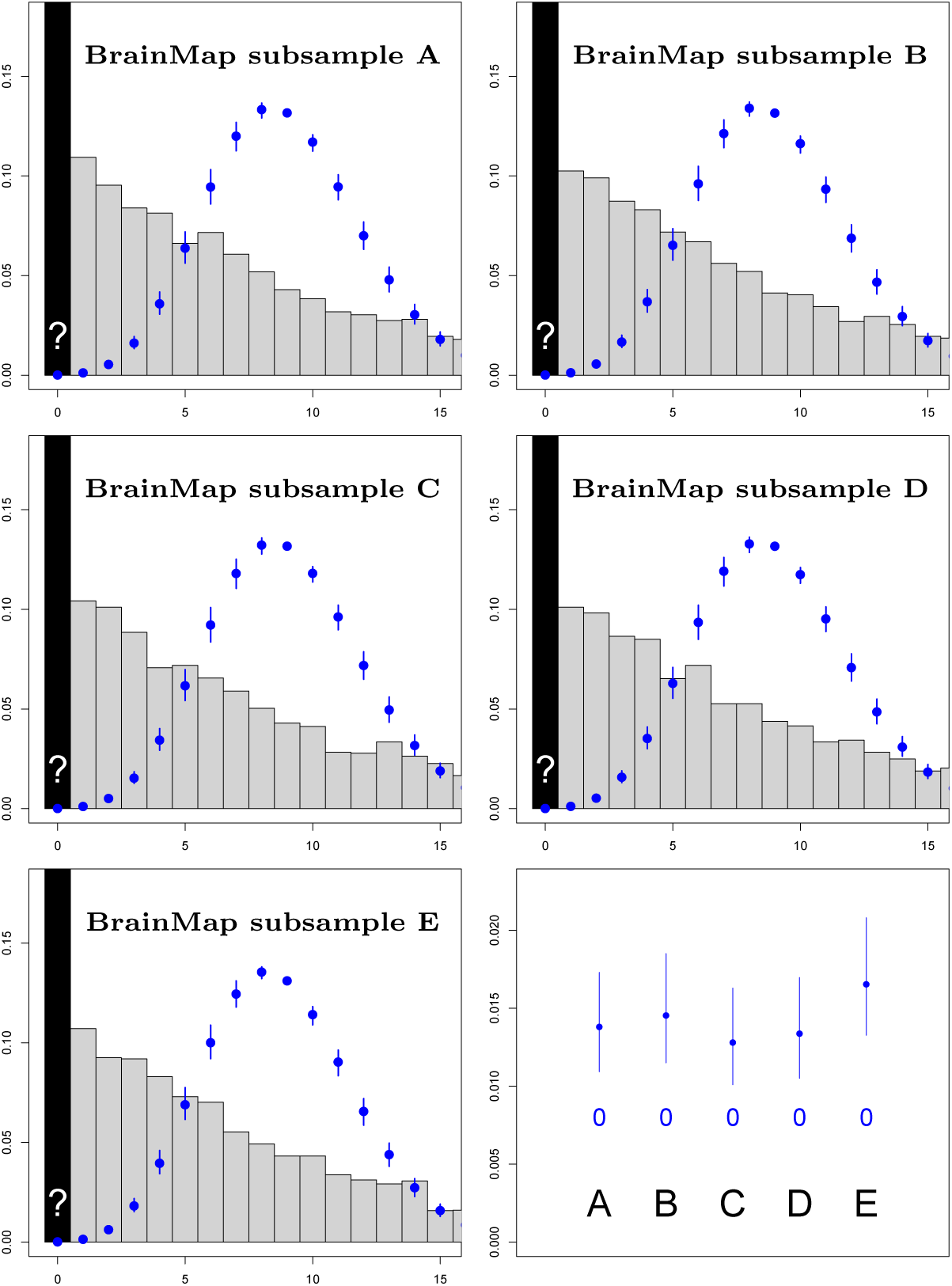
BrainMap results for 5 random samples using the zero-truncated Poisson distribution. The first 5 plots show observed count data (gray bars) with fit of full (non-truncated) distribution based on zero-truncated data, including the estimate of *p*_0_ (over black bar). Final plot shows estimates of *p*_z_, prevalence of file drawer experiments for every 100 experiments observed. All fitted values include 95% bootstrap confidence intervals. The Poisson model provides a poor fit to all 5 subsamples.

## C Negative Binomial and Delaporte parameter estimates

In this section, we present the parameter estimates obtained from the analysis of BrainMap subsamples A-E with the simple (without covariates) zero-truncated Negative Binomial and Delaporte models. The parameter estimates are listed in Table 8.

**Table 8:** Scalar parameter estimates obtained when fitting the simple zero-truncated Negative Binomial and Delaporte models to BrainMap subsamples A-E.

| Subsample | Negative Binomial |  | Delaporte |  |  |
| --- | --- | --- | --- | --- | --- |
| | $\mu$ | $\phi$ | $\mu$ | $\sigma$ | $\nu$ |
| A | 7.95 | 0.96 | 8.05 | 0.96 | 0.014 |
| B | 7.95 | 0.92 | 8.24 | 0.98 | 0.060 |
| C | 8.04 | 0.95 | 8.28 | 0.98 | 0.043 |
| D | 8.03 | 0.93 | 8.35 | 1.01 | 0.070 |
| E | 7.82 | 0.92 | 8.06 | 0.96 | 0.048 |

## Footnotes

1 1RRID:SCR 003069

2 Since we expect the power to scale with the square root of the sample size.

3 RRID:SCR 001905

4 RRID:SCR 002823

5 RRID:SCR 006798

